# Categorical encoding of voice in human superior temporal cortex

**DOI:** 10.1101/2021.11.23.469682

**Authors:** Kyle Rupp, Jasmine Hect, Madison Remick, Avniel Ghuman, Bharath Chandrasekaran, Lori L. Holt, Taylor J. Abel

## Abstract

The ability to recognize abstract features of voice during auditory perception is a complex, yet poorly understood, feat of human audition. For the listener, this occurs in near-automatic fasion to seamlessly extract complex cues from a highly variable auditory signal. Voice perception depends on specialized regions of auditory cortex, including superior temporal gyrus (STG) and superior temporal sulcus (STS). However, the nature of voice encoding at the cortical level remains poorly understoood. We leverage intracerebral recordings across human auditory cortex during presentation of voice and non-voice acoustic stimuli to examine voice encoding in auditory cortex, in eight patient-participants undergoing epilepsy surgery evaluation. We show that voice-selectivity increases along the auditory hierarchy from supratemporal plane (STP) to the STG and STS. Results show accurate decoding of vocalizations from human auditory cortical activity even in the complete absence of linguistic content. These findings show an early, less-selective temporal window of neural activity in the STG and STS followed by a sustained, strongly voice-selective window. We then developed encoding models that demonstrate divergence in the encoding of acoustic features along the auditory hierarchy, wherein STG/STS responses were best explained by voice category as opposed to the acoustic features of voice stimuli. This is in contrast to neural activity recorded from STP, in which responses were accounted for by acoustic features. These findings support a model of voice perception that engages categorical encoding mechanisms within STG and STS.

**Significance Statement:** Voice perception occurs via specialized networks in higher order auditory cortex, yet how voice features are encoded remains a central unanswered question. With human intracerebral recordings of auditory cortex, we provide evidence for categorical encoding of voice in STG and STS and occurs in the absence of linguistic content. This selectivity strengthens after an initial onset response and cannot be explained by simple acoustic features. Together, these data support the existence of sites within STG and STS that are specialized for voice perception.

## Introduction

Vocalizations are a crucial social signal and a fundamental driver of human and animal behavior. Humans and other animals can easily distinguish con-specific vocalizations from other complex sounds in their acoustic environment (1). With normal voice perception, one can quickly and accurately deduce gender, size, age, emotional state, and intentions purely from vocalization content (2). These voice recognition abilities begin to develop prenatally (3), precede development of linguistic abilities (4) and are formed through the processing of acoustic and paralinguistic aspects of voice (5). How humans are able to rapidly extract such rich information from vocalizations remains a central unanswered question.

Neuroimaging studies have identified regions of auditory cortex theorized to mediate voice processing which demonstrate robust BOLD response when listening to voice stimuli, including superior temporal sulcus (STS) and superior temporal gyrus (STG), collectively referred to as temporal voice areas (6–11). Recent neuroimaging work suggests that voice selectivity of STS and STG contributes to voice perception across primate species (1, 12–14). These regions exhibit robust connectivity with auditory regions in the supratemporal plane (STP), including Heschl’s gyrus, and higher order association cortices implicated in voice perception and voice identity recognition (15–18). Bilateral STS exhibits voice selective responses, although some studies suggest hemispheric asymmetry in STS anatomic structure and function (9, 19). Current understanding of regional voice selectivitity and temporal dynamics of these responses in auditory cortex is driven primarily by neuroimaging studies, therefore studies employing methods on physiologic timescales are needed to further examine the contribution of STS and STG to voice perception.

Whether activity in putative voice-selective areas of STG and STS are actually driven by voice and not the acoustic or linguistic properties of speech remains under debate (12, 20). A speech-driven model of voice coding is supported by some neuroimaging studies (7, 18, 20–23). Extant behavioral work also suggests voice perception may rely heavily on linguistic content (24). However, other studies have shown voice selectivity persists when controlling for the unique acoustic properties of voice stimuli (11). It remains unknown to what extent processing of complex auditory stimuli, such as speech, music, or naturally occurring environmental sounds, rely on shared or unique neural mechanisms (23, 25, 26) and how these mechanisms are organized across the auditory cortical hierarchy (27–30). Specific features driving neural encoding of voice and the timing and organization of this coding will also be advanced by approaches with greater temporal resolution.

To understand the cortical representation of voice at physiologic timescales, we measured local neural activity directly from the STS, STG, and surrounding auditory cortex in patient participants undergoing clinical intracerebral recordings as part of epilepsy surgery (31). To date, evidence for voice coding in human auditory cortex has largely been supported by fMRI studies. The low temporal resolution of fMRI limits interpretation of temporal dynamics of these responses, given physiologic delay and low-pass filtered nature of BOLD responses compared to peak spike frequency (32, 33). Here we leverage intracerebral recordings that uniquely allow direct electrophysiological measurements from sulcal banks, such as the STS and Heschl’s gyrus. We combined this recording technique with decoding and encoding modeling approaches to measure voice selectivity across the audiotry hierarchy and test the hypotheses that vocalizations are represented categorically in STG and STS. Here we provide data in support of voice category discrimination in neural recordings within STG/STS, and describe the temporal dynamics of these responses across temporal voice selective areas.

## Results

We recorded neural data from eight patient-participants undergoing intracerebral recordings as part of routine epilepsy surgery evaluation. Recording sites in each participant included sites in both the STP, STG, and STS, including Heschl’s gyrus (HG). Each participant performed an auditory 1-back task of natural sounds stimuli adapted from either Norman-Haginere et al., 2015 (20) (Natural Sounds; NatS) or Belin et al., 2000 (6) (Voice Localizer; VL). Each of these stimulus sets include vocal and non-vocal sounds that can be used to assess vocal selectivity similar to previous studies (6, 34). To measure local neuronal response to auditory stimuli, broadband high gamma activity (35–37), or HGA, was extracted (Fig. 1B and C), and each channel’s auditory responsiveness was assessed using a 2-sample t-test that compared mean HGA between a post-stimulus onset window (0 to 500ms) and a silent baseline (−600 to -100ms relative to onset). Channels that exhibited a significant auditory response (p < .05, FDR-corrected) were included in subsequent analyses. This resulted in between 28-72 auditory-responsive channels per patient, for a total of 399 channels.

**Figure 1.**
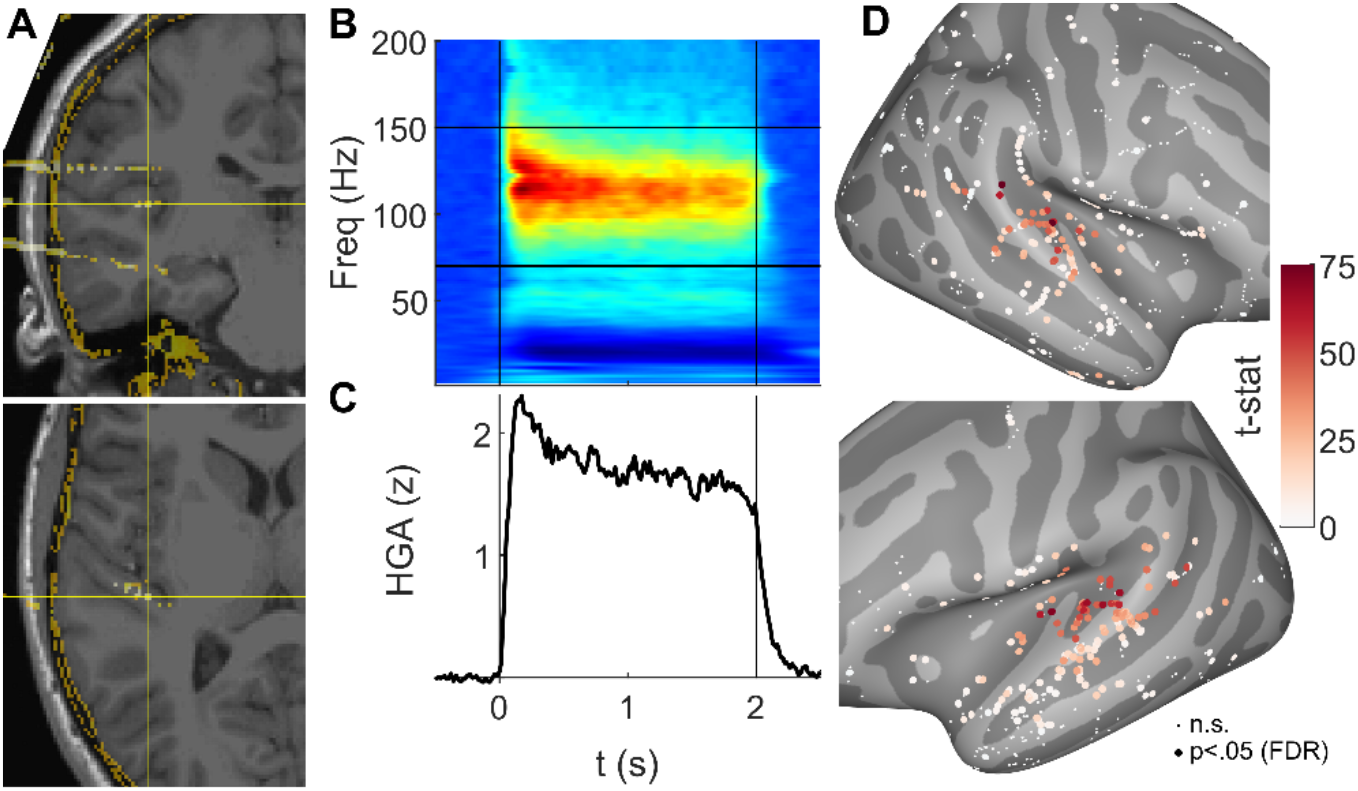
Auditory-evoked high gamma activity. (A) Example channel in left Heschl’s gyrus, patient P7. (B) Auditory-evoked spectral response averaged across all NatS stimuli in channel from (A). Vertical lines represent stimulus on- and offset, with horizontal lines demarcating frequency boundaries for broadband high gamma activity (HGA) at 70 and 150 Hz. (C) Mean HGA in the same channel. (D) Auditory responsiveness, quantified as the 2-sample t-value between mean HGA in 500ms pre- and post-stimulus onset windows. Small white dots represent channels with no auditory response, i.e., t-values that failed to reach significance (p < .05, FDR-corrected).

### Decoding Voice from Non-Voice Acoustic Stimuli

To establish the magnitude and temporal dynamics of vocal separability for the neural activity of each participant’s auditory responsive channels, we used HGA to decode between voice and non-vocal sounds. For full models (Fig. 2A), classification accuracy reached significance in each subject at p < .001 (permutation tests), with accuracy ranging from 82-93% for the VL stimulus set and 82-92% for NatS stimuli (see *Acoustic Stimuli*). To address whether separability was driven by encoding of linguistic information (i.e. speech) rather than voice, the full model decoding analysis was performed again with NatS data, excluding all speech stimuli (i.e., native and foreign speech, lyrical music), so that the vocal category contained only non-speech vocal sounds such as laughter, crying, and coughing. Accuracy ranged from 65-80% and remained statistically significant (p < .001 for 7 subjects, p < .01 for 1) (light blue bars, Fig. 2A).

**Figure 2.**
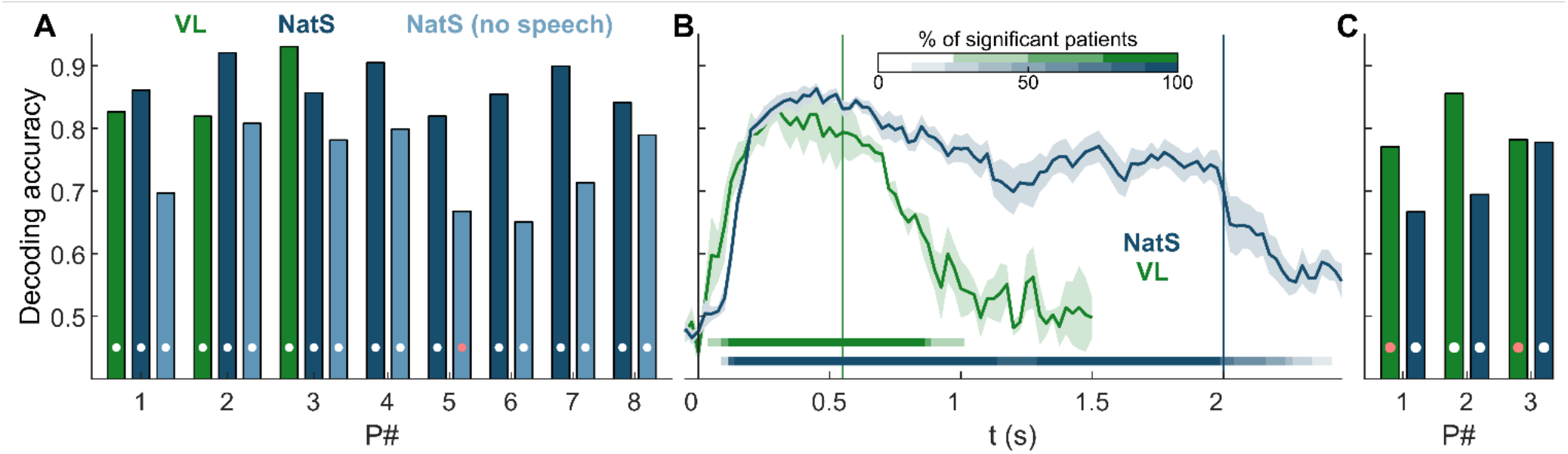
Decoding accuracy results. (A) Full model (i.e., all channels and time windows) decoding accuracy of vocal vs. non-vocal for each patient. Dark and light blue bars correspond to NatS results with speech stimuli included or excluded, respectively (e.g., light blue is non-speech human vocalizations vs. non-vocal auditory stimuli). Dots represent statistical significance (white: p < .001, red: p < .01, permutation tests). (B) Sliding window results. Vertical lines represent stimulus offset for the 2 tasks, with horizontal lines showing fraction of patients with statistically significant decoding in that window (p < .001, cluster-based permutation tests). (C) Cross-task decoding accuracy, with color indicating the training set (white: p < .001, red: p < .01, permutation tests).

Decoding accuracy time courses are shown in Fig. 2B, averaged across all patients for both NatS and VL. Across patients and stimulus sets, significant decoding emerged as early as 50ms (range of 50-150ms), with decoding accuracy falling below chance between 25ms before to 500ms after stimulus offset. Decoding accuracy trajectories were significant for the duration of the stimulus length for both NatS and VL, demonstrating that once vocal separability emerges, it persists for the duration of the sound.

Next, we examined the generalizability of our findings across different stimulus sets for the three subjects that performed both VL and NatS. Most notably, models trained on data from one stimulus set were able to decode vocal category membership on data from the other stimulus set (cross-task decoding, Fig. 2C), demonstrating similar HGA response properties despite completely distinct stimulus sets. Additionally, the sliding window accuracy profiles between VL and NatS were highly correlated within patient during the VL stimulus window (550ms; R = 0.80, 0.81, and 0.95 respectively).

### Distribution of Voice-Selective Sites

Next, to test the hypothesis that vocal separability is driven by activity in TVAs (STG/STS) we compared HGA across individual cortical recording sites in STP, comprised of HG and planum temporale (PT), versus STG/STS. Across all channels, STG/STS had smaller overall auditory HGA responses (p < 10^−5^) and decreased vocal – non-vocal (V-NV) separability (p < 10^−5^) relative to STP (Fig. 3A; rank-sum tests). We suspected that the latter finding, which ran counter to our a priori hypothesis, may have been driven by a significant difference in the proportion of channels showing significant V-NV separability (88% STP vs 52% STG/STS; p < 10^−5^, Fisher exact test). However, even when excluding non-significant channels, STG/STS did not exhibit greater V-NV separability than STP.

**Figure 3.**
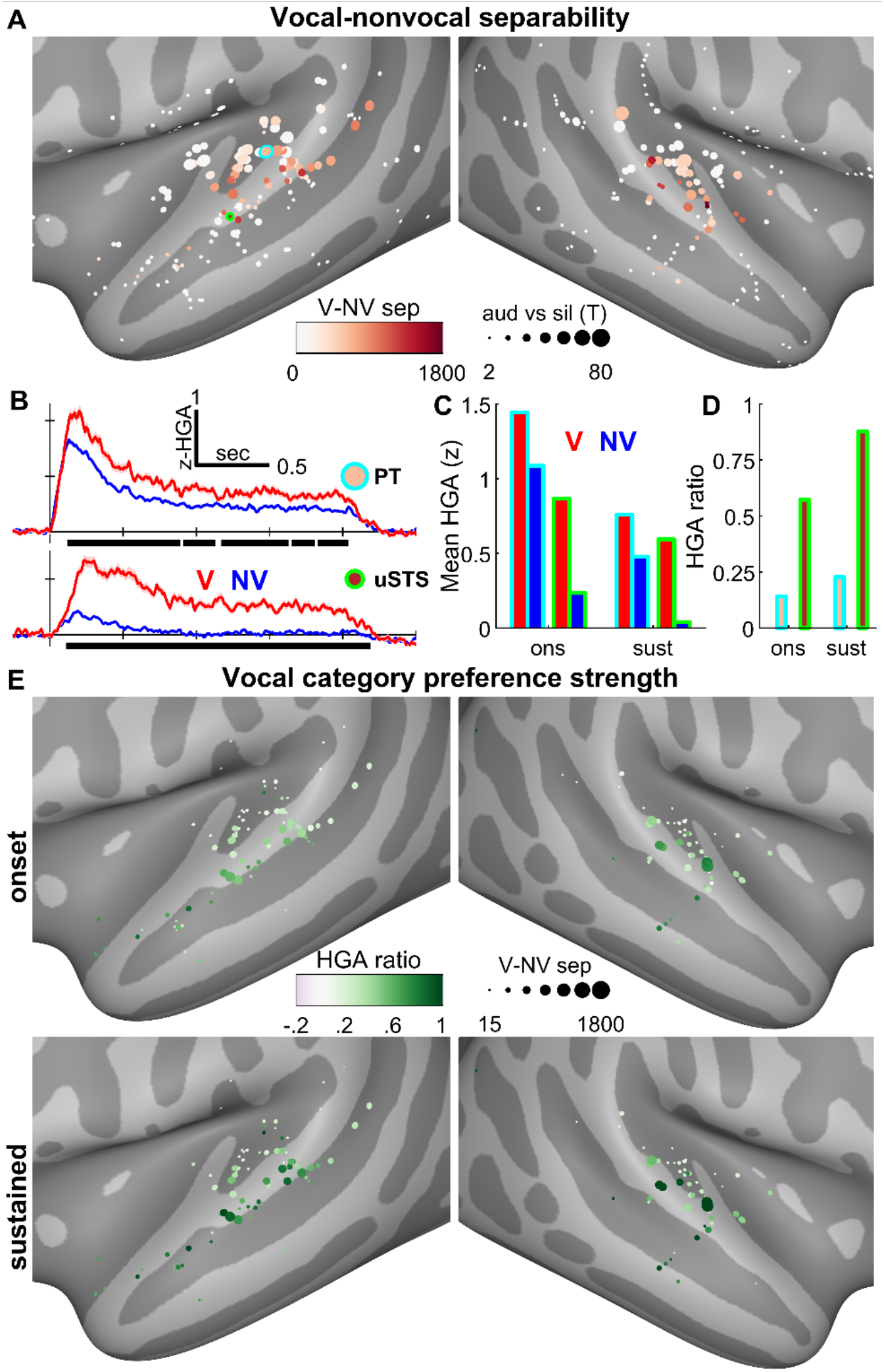
Single channel results. (A) HGA separability between vocal and non-vocal Natural Sounds stimuli, across all patients. Channel sizes are proportional to t-statistics comparing auditory response magnitude between 500ms pre- and post-stimulus onset windows, same as Fig. 1D. (B) HGA for 2 example channels located in planum temporale (upper panel) and near the STS/STG border (lower). Black bars show clusters of significantly different timepoints; V-NV separability (panels A, E) is the sum of all clusters for a given channel. Note that while both channels achieve V-NV separability throughout the duration of the stimulus, the magnitude of the non-vocal response differs between the 2 channels, with the NV response of the STG channel returning near baseline after the initial onset window. In contrast, the V response remains elevated in both onset and sustained windows, for both the PT and uSTS channels. (C) Mean HGA averaged across two different windows: onset (0-500ms) and sustained (500-2000ms). (D) The HGA ratio is calculated as the difference between vocal and non-vocal responses, relative to their sum. This metric, spanning from -1 to 1, describes a channel’s vocal category preference strength: a value near 1 (or -1) represents a channel that responds only to vocal (or non-vocal) stimuli, while a value of 0 represents equal HGA responses to both stimulus categories. (E) All channels with V-NV separability exhibit onset responses to both stimulus categories: in this early window, HGA ratios reveal that STG and STS (compared to STP) shows a slightly diminished response to non-vocal relative to vocal stimuli. During the sustained window, a strong preference for vocal stimuli emerges in STG and STS, while non-vocal responses return near silent baseline.

**Figure 4.**
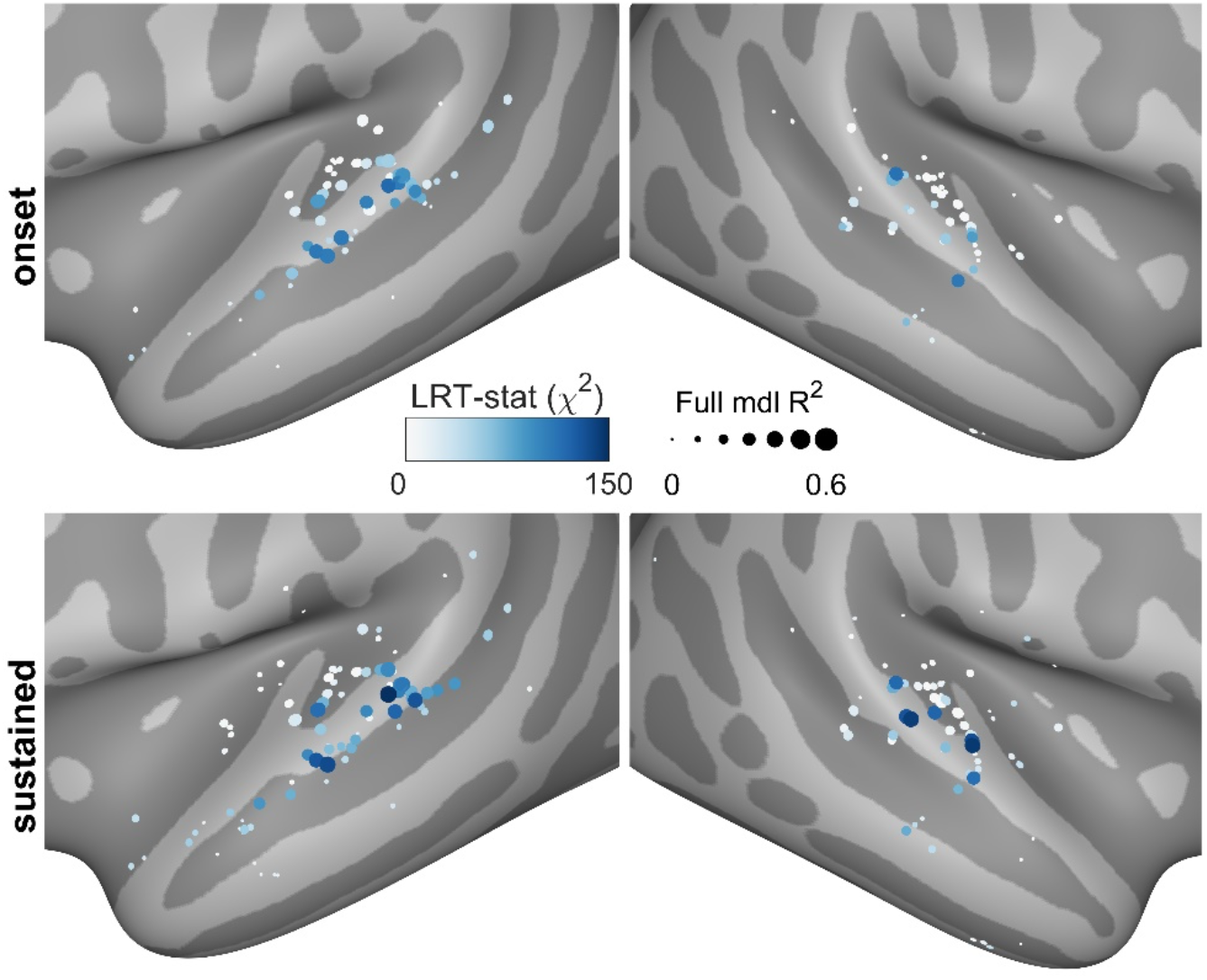
Encoding model results. Linear regression models demonstrate encoding of acoustic features in STP and category-like representations in STG and STS. Model inputs consisted of both low- and high-level acoustic features such as loudness, MFCCs, spectral flux, and relative formant ratios. Full models also included a binary feature indicating vocal category membership. Likelihood ratio test statistics compare this full model to a nested, acoustic-only model and thus describe the improvement conferred by V-NV class information. Well-fit channels in STP are modeled best by acoustic features throughout both the onset and sustained windows. Meanwhile, STG and STS channels also perform well and benefit from the addition of category-level information, with a slight skew toward the later sustained window.

The onset of V-NV separability was also estimated in STP and STG/STS. This onset is highly sensitive to the response strength, i.e., channels with poor signal-to-noise might show later separability onsets due to noise contamination. Therefore, we calculated the median onset time for only the 50% most separable channels in a given ROI. This resulted in median separability onsets of 130ms in STP and 150ms in STG/STS.

Next, we investigated the voice preference strength across the auditory cortical hierarchy. V-NV separability merely describes how distinguishable vocal responses are from non-vocal responses. A channel may be separable if both V and NV responses are elevated above baseline to differing degrees, as is the case with the PT channel in Fig. 3B, or if only V responses are elevated, demonstrated by the uSTS (upper bank of STS) channel. Responses exhibited 2 windows of distinct activity, consisting of an onset (0-500ms) and a sustained response (500-2000ms). The mean HGA response within these windows and across stimulus categories is shown for the 2 example channels in Fig. 3C. The category preference strength was then defined as the difference between mean V and NV responses, relative to their sum (Fig. 3D). Responses that are more exclusively confined to only V stimuli exhibit HGA ratios closer to 1, while ratios close to 0 represent much weaker category preference. Fig. 3D confirms the apparent trend in panel B: the cyan PT channel displays a weak category preference (HGA ratio < 0.25) throughout the stimulus duration, while the green uSTS channel transitions from a moderate to a strong category preference between the onset and sustained window.

Fig. 3E shows HGA ratio results across all separable channels (those with non-zero V-NV separability), revealing that the trend from Fig. 3B generalizes across channels. Specifically, across the auditory hierarchy, V-NV separability in the onset window is broadly driven by a weak category preference, strengthening to a stronger preference in the sustained window (p < 10^−5^, sign-rank test). Furthermore, STG/STS channels displayed a stronger vocal category preference relative to STP channels in both windows (onset: p=1.4×10^−6^, sustained: p=4.7×10^−6^, rank-sum tests). Taken together, these results suggest that STG/STS demonstrates a strong category preference that emerges only after an initial onset window during which all natural stimuli are processed.

### Voice Feature Encoding Demonstrates Category-Level Representation of Voice in the STS

Finally, we investigated whether categorical voice responses could be explained by lower-level acoustic processing. To this end, we used the audio-processing software OpenSMILE to extract acoustic features of varying complexity (38, 39). Specifically, we used the functionals feature set, which produces statistical summaries (e.g., mean, standard deviation, peak rate) of acoustic features (e.g., loudness, formants, mel-frequency cepstral coefficients, jitter, shimmer) for each stimulus. An additional binary feature was included to indicate vocal category membership. Encoding models were built to predict onset and sustained mean HGA (as in Fig. 3C) for each V-NV separable channel in auditory cortex using this feature space. Full encoding models used the full feature space, while nested encoding models were built without the categorical voice feature.

Two metrics were derived from this analysis that, when taken together, provide insight into a channel’s encoding properties. First, the percent of variance explained (R^2^) by the full model describes how well the input features explain the response magnitudes for a given channel. Second, the likelihood ratio test compares the nested to the full model and provides an estimate of the added value conferred by the introduction of the vocal category membership feature. Under the null hypothesis that both models fit the data equally well, this ratio is chi-square distributed.

Among encoding models with significant R^2^ values (p < .001, permutation tests), two qualitatively different types of responses emerged in the STP and STG/STS. The first group, clustered in STP, represents auditory feature encoding and is characterized by a combination of large R^2^ and low χ^2^ values. These channels are well-explained by encoding models in both the onset and sustained windows but show minimal improvement in model performance when a vocal category feature is added. The second group of channels, clustered in lateral STG and STS, demonstrates categorical encoding properties by showing substantial model improvement (large χ^2^) with the addition of categorical voice information (as well as large R^2^ values). ROI analysis confirms that χ^2^ values were significantly larger in STG/STS compared to STP in both the onset (p = .017) and sustained windows (p = .013, rank-sum tests). R^2^ values were not significantly different between regions in either window.

## Discussion

The human STS has long been associated with vocal category preference, but the exact computations that occur in the STS remain debated (20, 24, 40). One hypothesis is that, similar to face processing, temporal voice regions perform a voice detection gating function in which incoming auditory stimuli are categorized as vocal or non-vocal prior to higher level vocal processing (41–43). If such a model were correct, one would expect category-level encoding of vocal stimuli in temporal voice regions. Our results demonstrate cortical sites in the STG and STS, the putative sites of TVAs, have strong vocal category preference using two distinct voice localizer tasks. Notably, the data show that voice category preference strengthens along the auditory cortical hierarchy from STP to STG/STS in a temporally dynamic fashion, with an initial less-specific onset response followed by a sustained response with pronounced category preference in STG/STS. Importantly, our results also demonstrate that higher order voice preferring sites in the STG/STS are driven most robustly by voice category, rather than lower-level acoustic features. In contrast, voice-sensitive sites in the STP were driven primarily by acoustic rather than voice category features.

Whether voice category selectivity in human auditory cortex is driven by low-level acoustic features or whether neural response selectivity actually reflects more abstract representation of voice category remains an open question (12, 20, 24); some have suggested that voice specialization actually reflects specialization for speech (20, 21, 24, 44). First, our results show that while speech information plays a role in vocal neural coding, vocal decoding can occur even in the absence of speech. Second, our encoding model results demonstrate that in the STG/STS voice category plays a far more influential than low-level acoustic features. Thus, our results support a model in which there is a gradient of voice category selectivity across the auditory hierarchy, with lower-level acoustic features playing the most important role in supratemporal plane, and strong voice category selectivity emerging in bilateral STG/STS.

This study also sheds light on critical temporal dynamics of voice processing across the auditory system. Previous HD-EEG work reported an N200 signal distinguishing vocal from non-vocal stimuli with an onset at 164ms (45). Meanwhile, a subsequent MEG study showed a dissociation in the neural activity of vocal and non-vocal activity starting at 150ms (46). These studies both inferred a similar localization of this effect: the MEG study showed maximal dissociation around bilateral STS, while the HD-EEG study also proposed a similar anatomical locus. In agreement with these findings, we observed an onset of V-NV separability around 150ms in STG/STS channels. Interestingly, our decoding results revealed a slightly earlier onset across auditory cortex, starting between 50-150ms. These results may be due to decoding models exhibiting higher sensitivity to early weak separability, given the inclusion of multiple channels simultaneously.

While we found that separability between vocal and non-vocal responses is maximal in STP, several response characteristics suggest that voice selectivity is localized in STG/STS. Vocal category preference strength and categorical voice encoding are both stronger in this region, particularly during the sustained window following onset responses. In contrast, STP sites that display V-NV separability show a weak category preference strength, possibly related to acoustic feature encoding rather than true category specificity. This explanation is supported by the finding that responses in this region encode acoustic but not categorical features. Additionally, our results support the idea of dynamic selectivity (47) in the STS (i.e. there are two distinct phases of selectivity), whereby vocal category preference strength evolves from weak during the onset window to strong during the sustained response.

The NatS stimulus set was not designed as a voice localizer and thus possesses a lower proportion of non-speech vocal stimuli. To ensure this stimulus set functioned similarly as a temporal voice area localizer to VL, we performed a cross-decoding analysis between the natural sounds and voice localizer paradigms, which show that responses to vocal versus non-vocal sounds are similar across these separate stimulus sets (Fig 2C). This stands in contrast to functional neuroimaging work that used the NatS stimulus set to show that temporal voice regions may not exist (14). A recent study using artificially-generated sounds demonstrated that temporal voice regions may encode vocal perceptual quality, i.e., the extent to which a sound is voice-like (48). Since the environmental sounds stimuli do not sufficiently sample across this perceptual continuum, the current data are unable to shed light on this possibility directly. However, a weak (STP) versus strong (STG/STS) category preference strength could reflect encoding of acoustic and perceptual features respectively.

We demonstrate dynamic category-driven encoding of voice in the human STG/STS. Further, with the spatiotemporal resolution of intracerebral recordings, our results demonstrate a gradient of selectivity across auditory processing regions with distinct temporal dynamics underlying different aspects of voice processing. Taken together, our findings support a voice gating mechanism of voice coding by temporal voice regions.

## Materials and Methods

### Participants and electrode implantation

sEEG recordings of the STS, STG, and STP (including HG) were performed in 8 neurosurgical patients with drug-resistant epilepsy as part of clinical evaluation for epilepsy surgery. Written informed consent and assent (for patients >14 yrs old) was obtained from all subjects. The research protocol was approved by the University of Pittsburgh Institutional Review Board.

All patients underwent preoperative neuropsychological evaluation. sEEG electrode implantation was performed as previously described (31). Briefly, Dixi Medical Microdeep® electrodes were used, with a diameter of 0.8mm, contact length of 2mm, and center-to-center spacing of 3.5mm. Electrodes contained between 8 and 18 contacts each.

### Data collection

Patients performed an auditory 1-back task using short clips of natural environmental sounds (see next section for more details). Audio was presented binaurally via Etymotic ER-3C earphones, with volume adjusted to a comfortable level for each patient separately before the start of the experiment(s). Inter-stimulus intervals were randomized uniformly between 1-2 s. Patients were instructed to indicate 1-back stimulus repeats using a button box (Response Time Box, v6). Repeats occurred on about 16% of all trials. Each stimulus was presented a minimum of 2 (VL) or 3 (NatS) times, though the 1-back task design resulted in some stimuli with more presentations (up to 4 for VL and 5 for NatS).

Neural data were recorded at 1kHz using the Ripple Grapevine Nomad system (model R02000) with a real-time notch filter applied at 60, 120, and 180Hz. For patients P3-P8, the audio signal was split using a commercial splitter with separate volume controls; for P1-P2, the audio was fed into a distribution amplifier (Rolls, model DA134). In both cases, the split audio signal was presented to patients and simultaneously recorded synchronously with neural data by the Ripple amplifier.

### Acoustic stimuli

Two different types of stimuli were used in separate experiments, which we refer to as Voice Localizer (VL) and Natural Sounds (NatS). VL stimuli were modified versions of the stimuli used in the Belin voice localizer (6). These original stimuli were designed for fMRI experiments and consisted of 8 s clips, with each clip containing a series of either vocal (e.g., speech, laughter, coughing) or non-vocal (e.g., mechanical, music, animal vocalizations) sounds only. These stimuli were adapted to capitalize on the temporal resolution afforded by intracranial research: PRAAT silence-based segmentation was used to extract and save individual sounds from each clip (49). Sounds with duration shorter than 550 ms were discarded; all other sounds were shortened to this duration, linear-ramped on and off by 50 ms, and rms-normalized. This procedure generated 80 non-vocal and 72 vocal sounds; to ensure balanced classes, only the first 72 non-vocal sounds were selected.

NatS stimuli were the same as those originally used by Norman-Haignere et al. (20). Each of the 165 sounds are 2s in duration and belong to one of 11 categories, defined in the original study, which we grouped into superordinate categories of vocal and non-vocal sounds. Vocal categories consisted of English and foreign speech, human vocalizations, and lyrical music. Similar to VL, non-vocal sounds were more varied and included categories such as mechanical sounds, non-lyrical music, and animal vocalizations.

Importantly, the NatS stimulus set contained human non-speech vocal sounds that might not activate voice-selective regions of cortex. Specifically, crowd-generated cheering and laughter may be categorically different from vocal sounds generated by individuals. Furthermore, following the heuristic outlined by Belin et al. (6), we excluded sounds without vocal fold vibrations, namely breathing and whistling. Based on these two considerations, we reclassified four NatS stimuli from the vocal to the non-vocal category.

### Data preprocessing

A common average reference (CAR) filter was used to remove noise common across channels. Voltages were epoched by extracting a window of 1000ms before stimulus onset to 1450ms after offset in the case of VL, or 1000ms after offset for NatS. Each channel was then normalized relative to the pre-stimulus period across all trials. All channels whose centroid was further than 3mm from the closest cortical vertex (either the pial surface or the grey-white matter boundary) were excluded.

To estimate broadband high-gamma activity (HGA), epoched data was forward- and reverse-filtered using a bank of 8 bandpass Butterworth filters (6th order), with log-spaced center frequencies (70-150Hz) and bandwidths (16-64Hz). The analytic signal amplitude was extracted using the Hilbert transform. Each band was then normalized relative to a common baseline across all trials; in estimating the mean and standard deviation for normalization, the earliest 100 ms of baseline were discarded due to edge effects, and the 100 ms immediately preceding stimulus onset were discarded to prevent any contamination from low-latency responses. HGA was then calculated as the mean across these 8 bands, which was down-sampled to 100 Hz and clipped to a window of 900 ms before onset to 900 ms after offset.

Auditory-responsive channels were identified using 2-sample t-tests comparing mean HGA between a 500 ms window immediately following stimulus onset to a baseline period defined as -600 to -100ms pre-onset. Only channels with p < .05 (FDR-corrected) were used in subsequent analysis. For patients that completed both VL and NatS, channels were labeled auditory-responsive if they surpassed this threshold in at least one of the 2 tasks. At this point, HGA was averaged across all presentations of a stimulus, which we refer to as a stimulus response. Unless otherwise noted, stimulus responses (as opposed to single-trial responses) were used in all subsequent analysis.

### Decoding analysis

For each patient, decoding was performed via L1-regularized logistic regression using the MATLAB package *glmnet*. Input features consisted of mean HGA in 100ms windows, sliding every 50 ms. Relative to stimulus onset, window centers spanned from -50 to 1500 ms for VL and -50 to 2450 ms for NatS. Full models that included all channels and time windows, as well as sliding models that used single windows, were built.

In addition to regularization, cross validation (5-fold) was used to prevent overfitting and explore generalizability. Within each cross-validation fold, input features in the training and testing sets were z-scored relative to the training set. Additionally, each observation was weighted by the inverse of its class prevalence during the training phase, to prevent models from biasing toward the most numerous class. This step was especially important for NatS, due to a large class imbalance (37 vocal, 128 non-vocal stimuli). Lastly, an inner-loop 10-fold cross validation was used to test 20 different regularization parameters; the parameter was selected based on the model that minimized the mean deviance across inner folds. Balanced decoding accuracies were reported, in which the within-class accuracy for vocal and non-vocal stimuli were calculated separately and then averaged. Non-speech vocal decoding (Fig. 2A, light blue bars) was performed using single-trial (as opposed to stimulus) responses, due to the scarcity of NSV exemplars in the NatS stimulus set. For cross-task decoding, input features were limited to the shorter stimulus duration of VL, i.e., only windows within the first 550 ms. A single model was built on all data in one task and then tested on all data in the other task.

Statistical significance was assessed for sliding window decoding via a permutation-based clustering approach (50). Briefly, V-NV labels were shuffled, and sliding window decoding was performed 1000 times. At each window, this generated separate accuracy null distributions from which critical values were drawn, identified as the upper 95th percentile. For each permutation, values that exceeded their window’s critical threshold were saved, and temporally adjacent values were summed to create a cluster mass. The max cluster mass for each permutation was stored, generating a null distribution of 1000 cluster masses (if a permutation contained no windows that exceeded threshold, the max cluster mass was set to 0). Finally, the same procedure was applied to the true (unshuffled) sliding window accuracies, and each resultant cluster mass was assigned a p-value equal to the proportion of null cluster masses that exceeded it.

### Single-channel analysis

Single channel V versus NV separability was estimated using a similar approach. However, rather than using decoding accuracy, time-varying 2-sample t-statistics were calculated on single-trial responses. V-NV separability (Fig. 3A) was quantified as the sum cluster mass, i.e., the sum of masses for all significant (p < .001) clusters for a given channel. The importance of using this metric (as opposed to the max cluster mass) can be appreciated in the upper panel of Fig. 3B: summing across this channel’s 5 separate clusters gives a more accurate description of the overall separability.

Across V-NV separable channels, HGA response profiles appeared to share consistent morphological characteristics, namely onset and sustained responses of varying magnitudes. The longer stimulus durations in the NatS stimulus set provides a better estimate of sustained response properties and is thus the focus of this analysis. We first averaged HGA across an onset window (initial 500ms following stimulus onset) and a sustained window (remainder of the stimulus length, 500-2000ms), and then calculated the mean HGA within V and NV stimuli. To investigate response characteristics to V versus NV stimuli, we then calculated the HGA ratio, defined as the difference in mean HGA between these stimulus categories, normalized to the sum of these responses. Normalization helps account for a potential signal-to-noise confound: if a given channel’s overall response is scaled, the V-NV difference will be amplified as well.

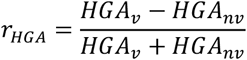

Assuming positive values for both HGA_v_ and HGA_nv_, this measure ranges between –1 and 1, with a value near 1 indicating a strong preference for vocal stimuli and a value near –1 indicating a strong preference for non-vocal stimuli. Among auditory-responsive channels that also demonstrated V-NV separability, a small fraction of them (7%) exhibited negative mean HGAs across NV stimuli, representing a NV-associated decrease in the HGA response relative to baseline. To constrain the HGA ratio between –1 and 1, these values were set to 0 before the calculation.

### Encoding models

By averaging across stimulus categories, the acoustic variability between individual stimuli has thus far been ignored. One possibility is that channels exhibiting a strong preference for vocal sounds are actually encoding lower-level acoustic properties that are inherently different between vocal and non-vocal sounds. To explore this possibility, encoding models were used to predict stimulus responses, i.e., mean HGA in both onset and sustained windows, for each channel that exhibited V-NV separability. Our approach closely mirrored the encoding model methods reported by Staib and Fruholz (48).

The OpenSMILE acoustic processing package, implemented in python, was used to extract the “functionals” set of 88 acoustic features for each NatS stimulus (38, 39). These features consist of statistical summaries (e.g., mean, standard deviation, percentiles) of acoustic features of varying complexity (e.g., loudness, mel-frequency cepstral coefficients, spectral flux, formant values). This feature space contained a high degree of collinearity between features; therefore, we used principal component analysis to reduce its dimensionality. The first n principal components that captured 99.99% of the variance in the original feature space were kept. Lastly, a categorical feature was added indicating vocal category membership. One stimulus (chopping food) was removed due to outliers in its acoustic features.

Linear regression encoding models were then built in one of two ways, corresponding to two relevant measures of interest. First, the overall model fit was calculated as the out-of-sample R^2^ value using leave-one-out cross-validation. Statistical significance of R^2^ values was assessed using permutation tests with 1000 permutations, in which rows of the feature matrix were shuffled before model building.

Second, the likelihood ratio test statistic was calculated between the full model and a nested version that excluded the vocal category feature. This statistic estimates the likelihood that the vocal category feature provides additional information beyond the acoustic features and is χ2 distributed under the null hypothesis. Log likelihoods for both full and nested models were attained from models trained on full stimulus sets.

## Funding

This work was funded by NIH NIDCD R01 DC013315 and NIH NIDCD R21 DC019217.

## Acknowledgments

We thank the participants, families, and epilepsy monitoring unit staff.

## Tables

**Table 1.** Participant epilepsy and behavioral characteristics.

|  | Summary | P1 | P2 | P3 | P4 | P5 | P6 | P7 | P8 |
| --- | --- | --- | --- | --- | --- | --- | --- | --- | --- |
| Age (years) | 15 (3) | 16 | 16 | 9 | 18 | 13 | 15 | 15 | 14 |
| Female, n (%) | 2 (25) | Male | Male | Male | Male | Female | Male | Female | Male |
| Right-handed, n (%) | 7 (87) | Left | Right | Right | Right | Right | Right | Right | Right |
| WASI-II score | 85 (16) | 76* | 62* | 67 | 95 | 94 | 81 | 102 | 104 |
| Left hemisphere language | 5 (63) | Left | Left | Left | Right | Bilateral | Left | Right | Left |
| Epilepsy onset (years) | 8 (5) | 1 | 0 | 8 | 12 | 3 | 10 | 13 | 13 |
| Seizure locus |  | Right parietal lobe | Right temporal lobe | Frontal lobe | Left frontotemporal, Heschl's gyrus | Right Amygdala | Mesial temporal lobe | Mesial temporal lobe | Left temporal gyrus |
| Epilepsy etiology |  | Unknown | Unknown | Autoimmune encephalitis | Autoimmune encephalitis | Mesial temporal sclerosis | Heterotopias and FCD | FCD | Pilocytic astrocytoma |
| Electrode contacts (auditory responsive / all) |  | 72 / 256 | 70 / 226 | 32 / 120 | 37 / 127 | 28 / 104 | 55 / 117 | 58 / 117 | 47 / 52 |
| NatSounds performance (% correct) | 85 (16) | 100 | 99 | 55.6 | 74.7 | 73.7 | 93.9 | 96 | 90.9 |
| NatSounds median RT (ms) | 1153 (346) | 929 | 1291 | 1064 | 1145 | 1912 | 818 | 1174 | 887 |
| VL performance (% correct) | 88 (15) | 100 | 71.4 | 91.1 | - | - | - | - | - |
| VL median RT (ms) | 837 (136) | 682 | 898 | 932 | - | - | - | - | - |

Summary statistics reported as mean (SD) unless otherwise specified; \* = WISC-V; FCD = focal cortical dysplasia

## Notes

### Competing Interest Statement

The authors have declared no competing interest.

